# *n*PoRe: *n*-Polymer Realigner for improved pileup variant calling

**DOI:** 10.1101/2022.02.15.480561

**Authors:** Tim Dunn, David Blaauw, Reetuparna Das, Satish Narayanasamy

**Affiliations:** University of Michigan, Ann Arbor, MI, USA

**Keywords:** germline variant calling, short tandem repeat (STR), copy number variant (CNV), nanopore sequencing, homopolymer, n-polymer, read alignment, variable gap penalty

## Abstract

Despite recent improvements in nanopore basecalling accuracy, germline variant calling of small insertions and deletions (INDELs) remains poor. Although precision and recall for single nucleotide polymorphisms (SNPs) now regularly exceeds 99.5%, INDEL recall at relatively high coverages (85×) remains below 80% for standard R9.4.1 flow cells [22, 23, 31]. Current nanopore variant callers work in two stages: an efficient pileup-based method identifies candidates of interest, and then a more expensive full-alignment model provides the final variant calls. Most false negative INDELs are lost during the first (pileup-based) step, particularly in low-complexity repeated regions. We show that read phasing and realignment can recover a significant portion of INDELs lost during this stage. In particular, we extend Needleman-Wunsch affine gap alignment by introducing new gap penalties for more accurately aligning repeated *n*-polymer sequences such as homopolymers (*n* = 1) and tandem repeats (2 ≤ *n* ≤ 6). On our dataset with 60.6× coverage, haplotype phasing improves INDEL recall in all evaluated high confidence regions from 63.76% to 70.66% and then nPoRe realignment improves it further to 73.04%, with no loss of precision.

## 1 INTRODUCTION

As long read technologies have matured and basecalling accuracy has increased to over 99%, their popularity has grown accordingly [2, 11]. Long reads are essential for spanning repetitive regions and unambiguously mapping reads. Last year, the first gapless human genome sequence was constructed by the T2T consortium by combining PacBio HiFi and ONT nanopore long reads [18]. Nanopore sequencing in particular has gained popularity due to its impressive read lengths, low cost, real-time results, and direct calling of base modifications [8, 26, 30].

The two current leading nanopore variant callers are Clair3 (developed by the HKUCS Bioinformatics Algorithm Lab) and PEPPER-Margin-DeepVariant (a collaboration between UCSC and Google Health, hereafter referred to as PEPPER) [23, 31]. Both tools have converged on a similar variant calling pipeline: basecalling, read alignment, pileup-based variant calling (using pileup summary statistics), read phasing, and full-alignment variant calling (using all read information).

Despite posting impressive F1 scores (≥ 0.995) for SNP calling, nanopore variant callers struggle with accurately identifying INDELs in low-complexity regions [14, 23, 31]. Most recent nanopore variant calling advances in this area have come from improvements in machine learning and data representation. For example, the move from prior work Clairvoyante [13] to Clair [14] involved *“an entirely different network architecture and learning tasks”*. Clair3 then split the model into a pileup caller to filter out the noise and a higher-dimensional full-alignment caller to make the more difficult decisions [31]. PEPPER examined sorting reads by haplotype and a new architecture, and DeepVariant explored numerous possible data representations for final calling [19, 23]. Orthogonally, we show that improved INDEL calling performance can be achieved through better read alignment by introducing novel gap penalties for homopolymers and tandem repeats, or “*n*-polymers”.

In order to maximize the accuracy of pileup-based variant calling, reads should be aligned such that actual mutations are always aligned to the same location, despite sequencing errors. We find that simply using traditional affine gap penalties is not ideal because gap penalties *G*_open_ and *G*_extend_ are static, regardless of context [5]. For example, although our dataset consisted of only 0.8% INDEL errors, homopolymers of length 10 contained an INDEL error 41.8% of the time. Without lowering the INDEL penalty in the context of repetitive sequences, common sequencing errors have an outsized impact on fine-grained read alignment, often at the expense of consistently aligning actual mutations.

Figure 1 demonstrates a specific example where static INDEL gap costs cause poor alignment concordancy in low-complexity regions. Reads are identical in the two pileups shown, only the alignments differ. In this example, two adjacent homopolymers are basecalled with inconsistent lengths. nPoRe recognizes that these two events were most likely independent, and separates them into two homopolymer length mis-calls/variants. In contrast, MiniMap2 merges two INDELs whenever possible, or aligns homopolymer length differences as SNPs when one homopolymer is lengthened and the other is shortened, resulting in inconsistent alignment. According to the truth VCF, the first homopolymer of all As had a single deletion. Looking at the third base in the coverage graphs in Figure 1, we can see that nPoRe placed a deletion here for a much larger fraction of reads than MiniMap2.

**Figure 1:**
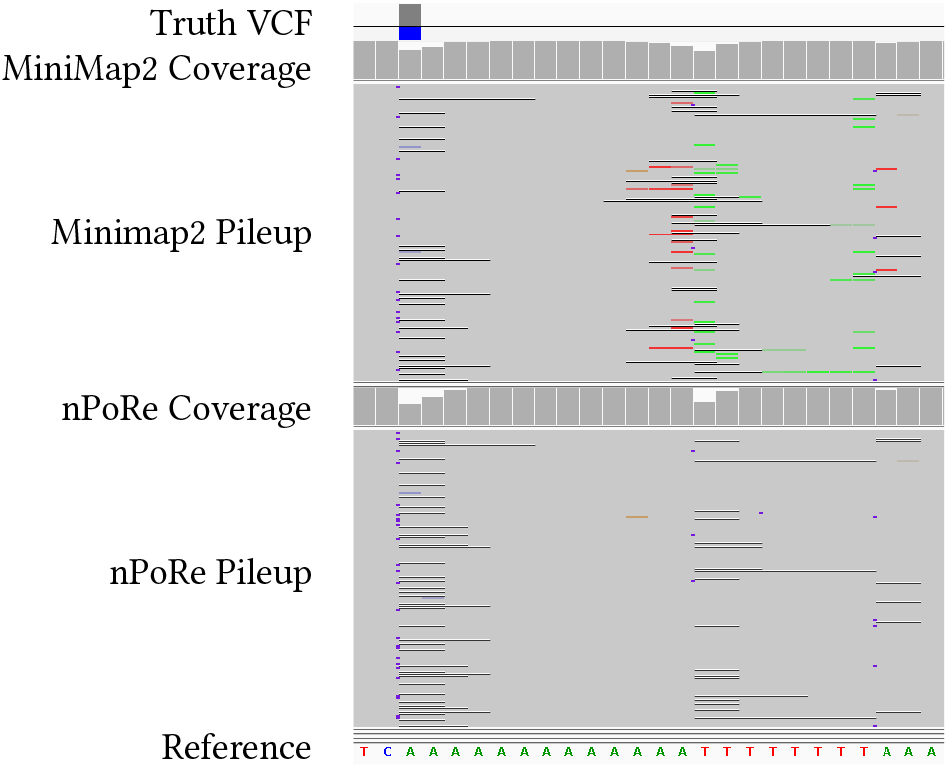
MiniMap2 and nPoRe aligned reads [21].

The likelihood of incorrectly basecalling an INDEL within a homopolymer increases significantly as homopolymer length increases. Figure 3a shows the confusion matrix for actual and base-called homopolymer lengths in our dataset. This same trend is visible for tandem repeats of longer length, though to a lesser extent (Figure 3b).

**Figure 2:**
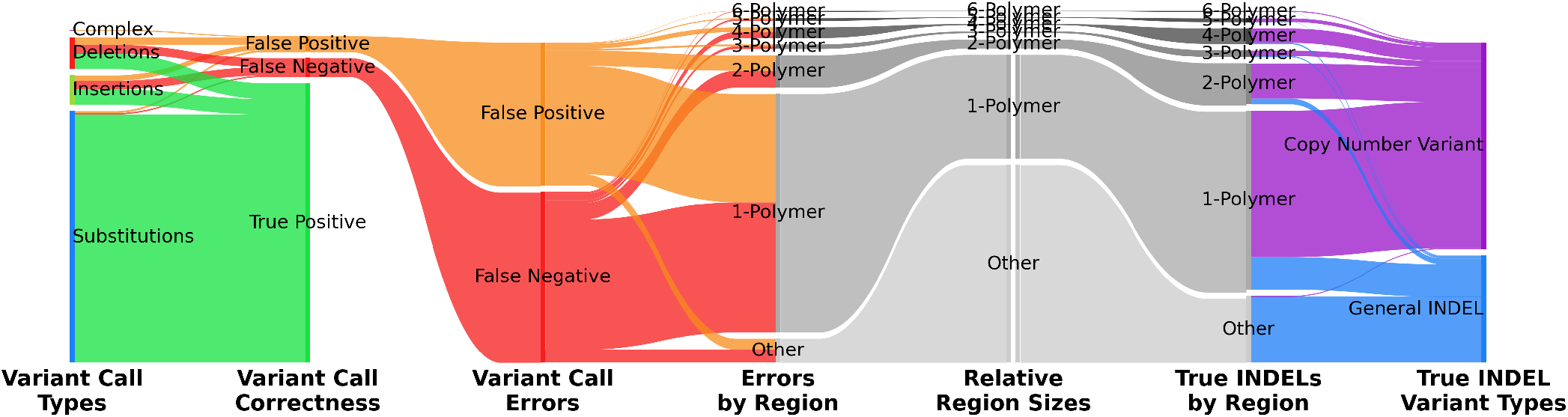
Sankey diagram demonstrating proportion of pileup-based variant calling errors (left) and true INDEL variants (right) contained within designated *n*-polymer (as defined in Section 3.2) regions of `chr20-22` in GM24385.

**Figure 3:**
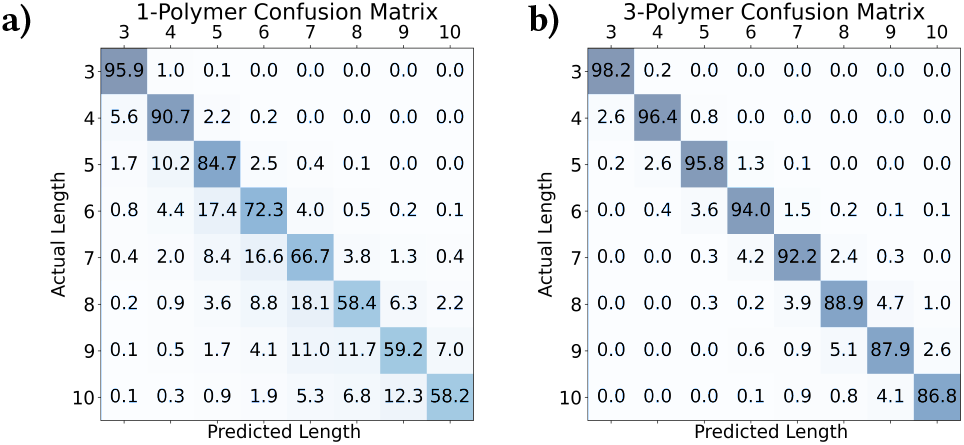
a) 1- and b) 3-polymer confusion matrices (in %)

Nanopore-based variant callers have historically struggled with INDELs, particularly with recall. Although Clair3 achieves 99.67% precision and 99.60% recall for SNP variant calling, it achieves only 90.86% precision and 64.73% recall for INDELs [31]. PEPPER v4 performs similarly, with 99.61% and 99.62% SNP precision and recall but just over 90% precision and 60% recall for INDELs [23]. The most recent evaluation available shows PEPPER v7 achieving 93% precision and 76% recall for INDELs, at 85× coverage [22].

Our own evaluation confirms these findings, and furthermore attributes the loss of INDEL recall to the first pileup-based variant calling step. Figures 4a and 4b show SNP and INDEL precision recall curves, respectively, for both Clair3’s pileup and full-alignment models. Note that although the more complex full-alignment model significantly improves precision, it cannot improve recall as dramatically; only variant calls and low-confidence reference calls from the previous pileup-based step are considered.

**Figure 4:**
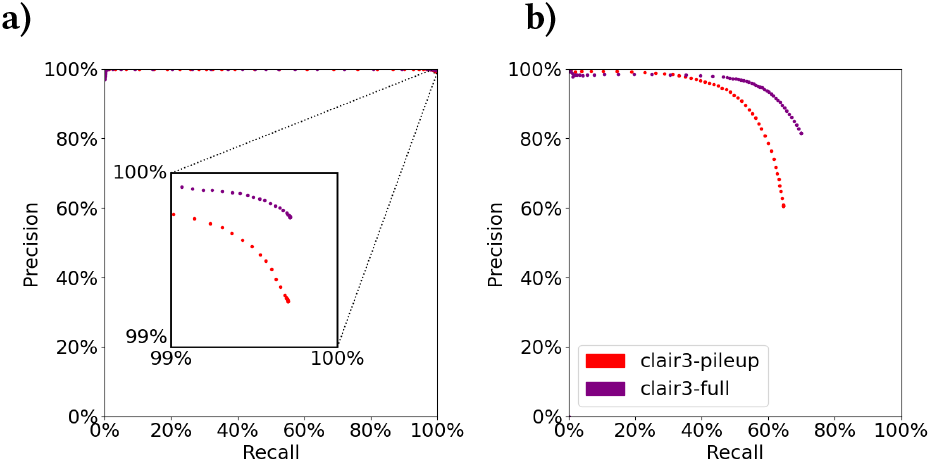
a) SNP and b) INDEL germline small variant calling accuracy of baseline clair3-pileup and clair3-full.

Figure 2 shows that although substitutions comprise 83.75% of the actual germline variants in our dataset, INDELs account for 92.36% of the pileup-based false negative and 80.79% of false positive variants. Of these errors, 92.29% occur within *n*-polymer regions, despite *n*-polymer regions covering just 37.07% of evaluated regions. By improving the alignment of reads in these small *n*-polymer regions, we can have a significant impact on overall variant calling accuracy.

INDEL mutations are over-represented in *n*-polymer regions as well (79.64% of all INDELs). This is because Copy Number Variants (CNVs) are a common form of mutation due to strand slippage during DNA replication. We define copy number variants as *n*-polymers (3+ exact copies of the same repeat unit), with a differing number of copies from the expected reference. For example, AAAA→AAAAA and ATATAT→ATAT meet this definition, but ATAT→ATATAT, AATAATAAAT→AATAAT, and ATATAT→ATATA do not. Despite our relatively strict definition of *n*-polymer copy number variants, however, 65.82% of all INDELs met this classification. nPoRe’s algorithm is directly designed to reduce alignment penalties for CNVs and improve alignment in such low-complexity regions.

Our work makes the following contributions:

- We identify the main source of nanopore germline small variant calling errors to be copy number variant false negatives in tandem repeat and homopolymer (“*n*-polymer”) regions during pileup-based variant calling
- We show that context-agnostic affine gap penalties do not accurately reflect the likelihood of nanopore CNV sequencing errors in *n*-polymer regions
- We extend Needleman-Wunsch affine gap alignment to include context-dependent gap penalties for more accurately aligning *n*-polymers
- We introduce “follow-banding” for efficient realignment We develop a VCF standardization script that ensures variants are reported in the same format as our realigner
- We show that haplotype phasing and nPoRe realignment significantly improve pileup-based variant calling accuracy

## 2 RELATED WORK

Variable gap penalties have been around for a long time. In 1995, Thompson first introduced per-position gap opening and extension penalties [29]. Since then, the sub-field of homologous protein sequence alignment has made extensive use of variable gap penalties (PIMA [25], FUGUE [24], and STRALIGN [4]) due to a high correlation between INDEL likelihood and the existence of protein secondary structures such as *α*-helices and *β*-strands. SSALN was the first to use empirically-determined penalty scores (an approach similar to our own) [20], and SALIGN greatly increased the flexibility of the gap penalty function, although with a corresponding increase in computation [15]. Unfortunately, all of these early works were designed to operate on short sequences and the numerous extensions of traditional Needleman-Wunsch alignment devised cannot be directly applied to *n*-polymer alignment.

Previous research projects explore proper alignment of Short Tandem Repeats (STRs), although most function as INDEL variant callers, rather than read realigners. Several newer works do focus on read realignment, however. ReviSTER is one such tool for revising mis-aligned/mapped reads through reference reconstruction with local assembly, though this is primarily helpful for improving mapping, not alignment. The Broad Institute has incorporated into their standardized analysis pipeline (Genome Analysis ToolKit, or “GATK”) an IndelRealigner, recognizing that INDELs are frequently mis-called as SNPs at read edges [7]. STR-realigner flags repeated regions and aligns them separately, allowing repeated traversal of STRs during alignment [9]. They find that this approach improves the consistency of read alignment in and near repeated regions, improving downstream variant calling. Despite this, no existing STR realigners are designed for long reads.

## 3 ALGORITHM

### 3.1 Overview

Because we have designed a read **realignment** algorithm, we trust the initial mapping of each read. Each read and its corresponding section of the reference genome are realigned, and a new traceback (alignment path) is computed. In other words, our solution simply adjusts the CIGAR string of each read within the input BAM file to better model the most likely mutations and sequencing errors in an effort to achieve greater concordancy between reads.

Our realignment algorithm is an extension of the Needleman-Wunsch algorithm for global alignment [17]. In addition to including known improvements such as an affine gap penalty and custom substitution penalty matrix [1], our algorithm allows the shortening and lengthening of homopolymers and tandem repeats (i.e. ACACAC→ACACACAC).

### 3.2 *n*-Polymer Repeats

The literature often categorizes sequences consisting of one repeated base as “homopolymers”, and repeated sequences of at least two bases as “tandem repeats” or “copolymers” [10, 23]. Short tandem repeats (STRs) are often defined as repeated units 2-6 bases in length, and are also known as “microsatellites” or “simple sequence repeats” (SSRs) [3]. Rather than treating these classifications separately for nPoRe, **we define an *n*-polymer to consist of at least 3 exact repeats of the same repeated sequence, where the repeat unit is of length 1 to 6 bases** (1 ≤ *n* ≤ 6). For example, homopolymers such as AAAAA (*n* = 1) and tandem repeats such as ACACACAC (*n* = 2) and TTGTTGTTG (*n* = 3) are *n*-polymers. Shorter or irregular repeated sequences such as AATTAATT and ACAACAACAC are not.

### 3.3 Penalty Functions

For each read, differences from the reference genome can be attributed to either sequencing errors or actual mutations. Regardless of origin (error or mutation), our goal is to align these reads to the reference in a manner that accurately captures the change that occurred. Existing aligners fail to do this by defining substitution and gap penalties based on estimated rather than measured rates of occurrence, and the algorithms do not account for common sequencing error modes such as tandem repeat length errors in nanopore sequencers. In contrast, we calculate penalties based on frequency measurements from the input BAM file. We define the penalty score for each difference (whether error or mutation) to be the negative log likelihood of that event occurring. As a result, finding the minimum-penalty alignment path is equivalent to finding the most likely set of errors and mutations that have occurred (assuming independence).

#### 3.3.1 Substitution Penalty Matrix

Figure 5 shows the calculation of substitution penalty matrix *P* from confusion matrix *C*_*P*_, using Equation 1. *ϵ* = 0.01 was included for numerical stability in the case that certain events were never observed. If we consider bases *x* = “*ACGT* “, then *P[i, j]* is the negative log probability that base *x[i]* was observed as base *x[j]*, either through a mutation or sequencing error:

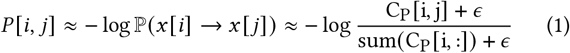

**Figure 5:**
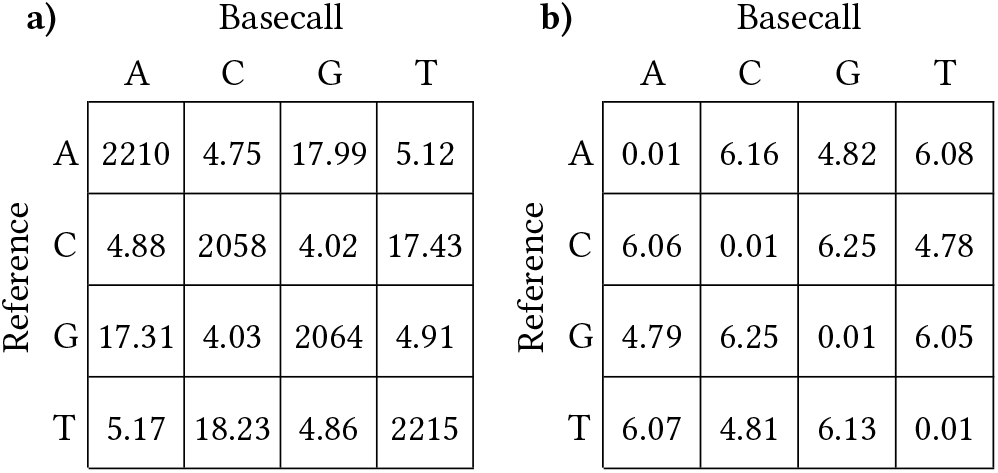
a) substitution confusion matrix *C*_*P*_, count in millions, and b) resulting penalty matrix *P*.

#### 3.3.2 Affine Gap Penalties

Confusion matrices for insertions (*C*_*I*_) and deletions (*C*_*D*_) were first generated by measuring the occurrence of small INDELs in the input BAM. Both matrices are 1D, since the expected INDEL length is always zero. Then, penalties were calculated by determining the negative log probability of each INDEL length *i* occurring: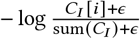. From these penalties, a best-fit gap opening penalty *G*_open_ of 5 and gap extension penalty *G*_extend_ of 1 was selected for both insertions and deletions [1].

#### 3.3.3 Tandem Repeat Penalty Matrix

First, confusion matrix *C*_*N*_ of shape 6 ×100 ×100 was generated by comparing expected and observed *n*-polymer lengths *l* (up to 100). For each *n*, or repeat unit size 1 − 6, a penalty matrix was calculated using the following equation, where *i* is the expected repeat length and *j* is the measured repeat length.

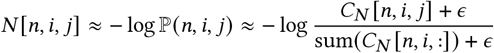

To improve penalty regularity, particularly for longer *n*-polymers where few examples were observed, the following two properties were enforced:

- shorter INDELs are more likely:

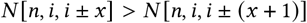
- same-length INDELs are more likely in longer *n*-polymers:

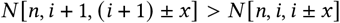

### 3.4 Reference Annotation

Reference annotations are used to track eligible *n*-polymers during alignment. For each possible *n*-polymer repeat unit length from *n* = 1 to *n*_max_, each reference position is annotated with *l*, the length or number of consecutive repeat units, and *idx*, the 0-based index of the current repeat unit (0 ≤ *idx < l*). Table 1 shows example annotations for a short reference sequence for *n* = 1 and *n* = 2. Recall from Section 3.2 that in order for a sub-sequence to be considered an *n*-polymer, the pattern must repeat exactly at least three times. Annotations may overlap, and non-zero annotations are only placed at the start of every *n*-polymer repeat unit.

**Table 1:**
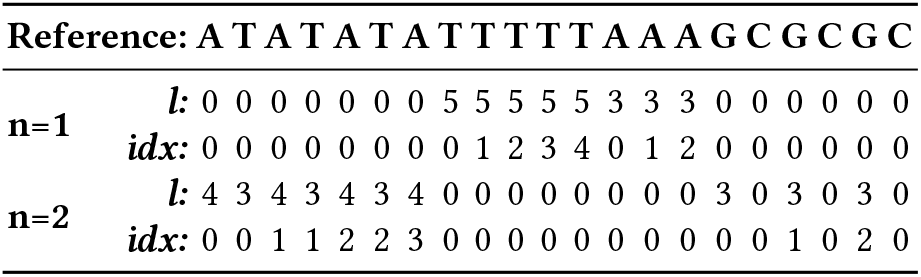
Example *n*-polymer reference annotations.

### 3.5 Alignment

Before aligning read *r* to reference *R*, the reference is annotated with *n*-polymer information as discussed in Section 3.4. Then, the five matrices *D, I, M, S*, and *L* are computed in lockstep one cell at a time, in that order. These matrices of size |*r*|× |*R*| represent the states Deleting, Inserting, Matching, Shortening *n*-polymers, and Lengthening *n*-polymers, respectively. For each cell, each matrix stores a tuple (*val, pred, run*) containing the accumulated penalty *val*ue, in addition to the *pred*ecessor matrix and consecutive movements (*run*) within that matrix for backtracking purposes.

Figure 6 demonstrates the cell dependency patterns and penalties in greater detail. For example, when computing cell *i, j* in *S*, the reference annotations *l* and *idx*, are first retrieved for each *n* for *R*[*j*]. All dependencies (marked with ^•^ for *S*) in Figure 6 are considered, and the minimum value of these dependencies’ cell values plus the associated penalties is calculated and stored in the result cell (marked with • for *S*). In other words, when looking at *S* [*i, j* ], for each *n*, we do:

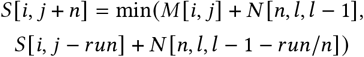

**Figure 6:**
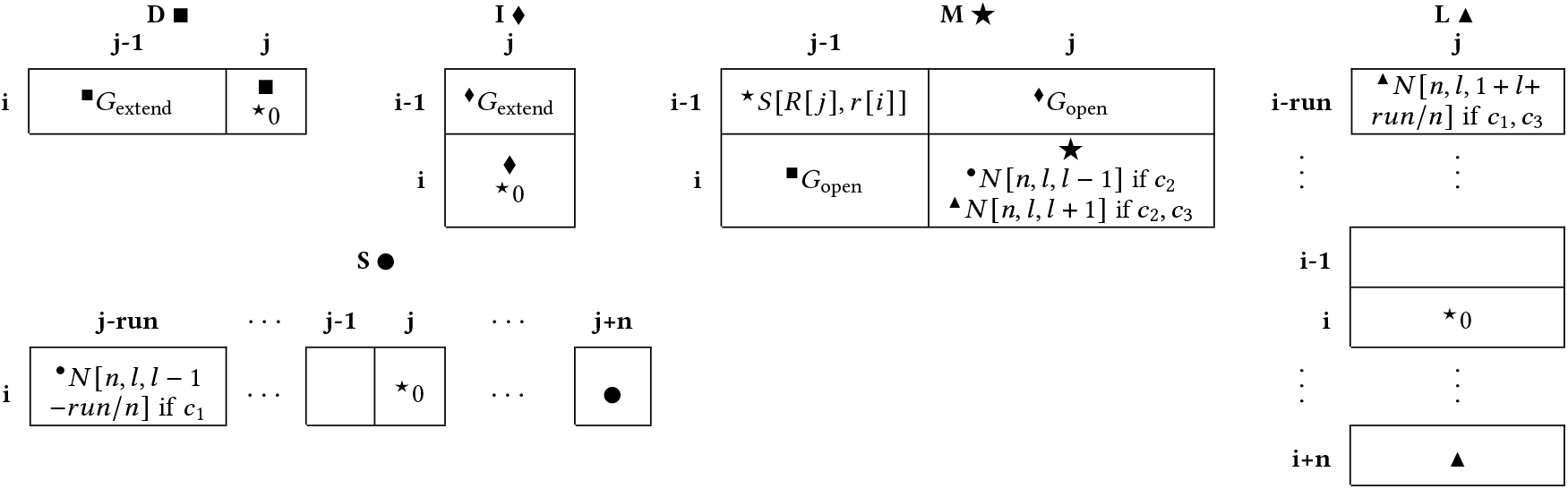
Realignment algorithm dependencies and penalties for computing cell *i, j* in the five matrices *D*■, *I* ♦, *M*⋆, *S*•, *L*▲, **which represent the states Deleting, Inserting, Matching, Shortening, and Lengthening, respectively. Computation for cell** *i, j* occurs in that order, and large symbols (e.g. ■) denote where the result is stored. Superscript symbols (e.g. ^■^) represent cell dependencies, and a penalty score accompanies it. Each result is the minimum value of all dependency cells plus their accompanying penalty scores. *n*-polymer shortening and lengthening is only allowed if certain conditions are met (*c*_1_, *c*_2_, *c*_3_), described further in Section 3.5.

These two movements correspond to starting to shorten a tandem repeat (state *M* →*S*), and continuing to shorten a tandem repeat (state *S* →*S*). All movements into matrices *S* and *L* such as these are only allowed conditionally based on reference annotations (described in greater detail in Section 3.6). Note that if matrices *S*• and *L*▲ are omitted (as well as all ^•^ and ^▲^ dependencies), this algorithm is equivalent to Needleman-Wunsch alignment with an affine gap penalty [17].

### 3.6 *n*-Polymer INDEL Conditions

Unlike matrices *D, I*, and *M*, the results for matrices *S* and *L* are stored several cells ahead of the current cell, and cell dependencies are only allowed conditionally based on the reference annotations. This ensures that matrices *S* and *L* only allow INDELs which change the copy number of tandem repeats. Here are the three conditions *c*_1_, *c*_2_, *c*_3_ used by our algorithm, and referenced in Figure 6:

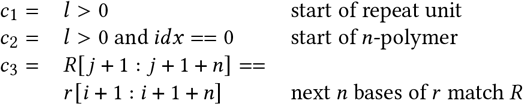

### 3.7 Backtracking

Traceback occurs entirely within matrix *M*, and relies on *pred*ecessor and *run* length information computed during the forward pass. The selected optimal alignment path is computed by Algorithm 1 and reported in the output BAM file in the form of a CIGAR string. CIGAR strings are composed of the symbols M, I, and D. Reference Matches, Insertions, and Deletions, correspond to diagonal, vertical, and horizontal movements in the alignment matrix, respectively. An example alignment is denoted by • in Figure 7a. After computation, the computed CIGAR string is collapsed (MMMDMIMM→3M1D1M1I2M).

**Figure 7:**
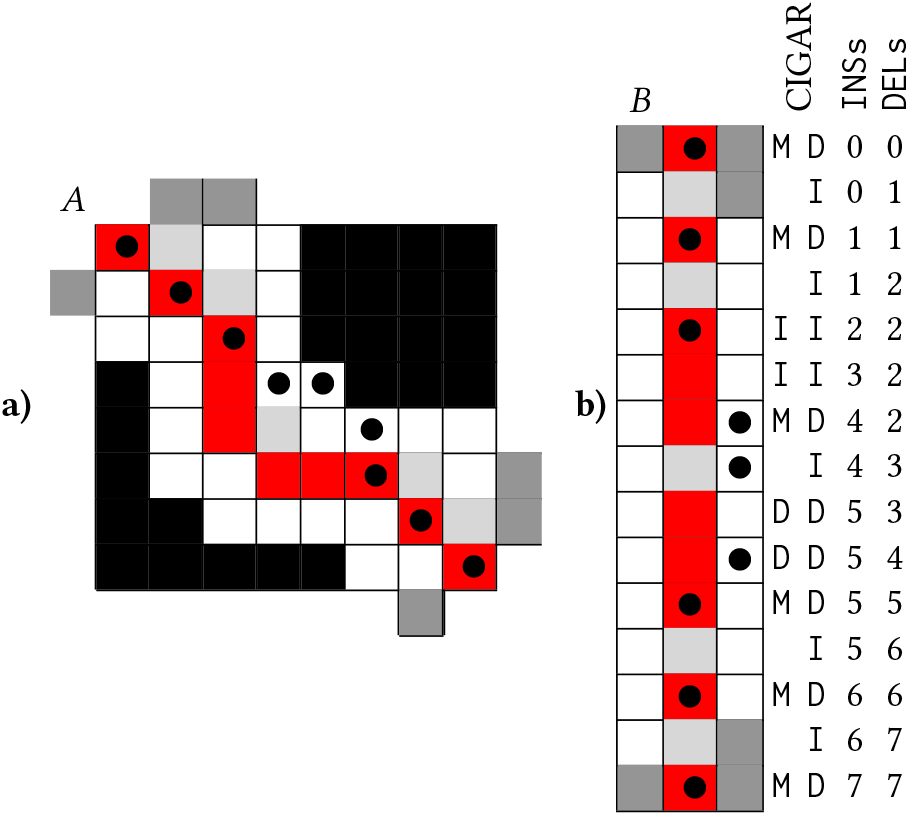
Follow banding matrix transformation *A* → *B*.

### 3.8 Follow-Banding

As mentioned previously, the primary goal of our read realignment algorithm is to more accurately model the mutations and sequencing errors in fine-grained alignment. Therefore, we can skip read mapping and trust the read start position reported by the previous aligner.

#### Algorithm 1 Backtracking algorithm

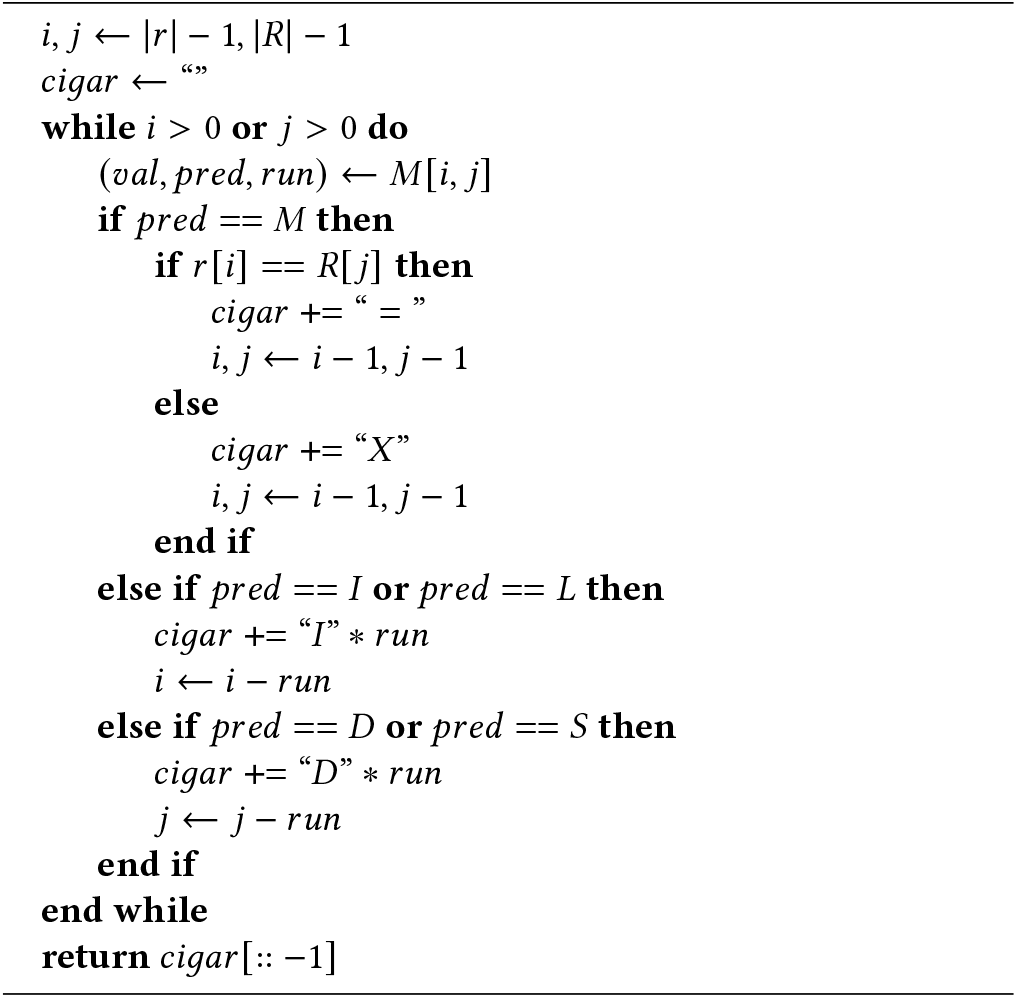

In addition to the mapping `POS`ition field, the SAM/BAM file format [6] includes `TLEN`, or the template (reference) length which corresponds to the aligned read. Thus, we can align read *r* to a small subsection of the reference *R* `[POS : POS TLEN]`. Barring any large INDELs, TLEN ≈ |*r*|, and we transform our alignment problem from *O* (|*R*| |*r*|) to *O*(| *r* |^2^).

Moreover, we can use the SAM/BAM file’s existing CIGAR string to simplify our alignment problem even further. Our optimal alignment will likely follow a path close to that of the original alignment. Figure 7a demonstrates how we can compute the alignment matrix (new optimal path denoted by •) in a narrow band *b* = 1 that follows the original alignment (red cells). Dark gray and black cells are not computed. Essentially, we use the CIGAR string to precompute the movement directions for adaptive banded alignment as proposed by Suzuki and Kasahara [27], instead of using a heuristic comparing penalty scores on the band’s edge.

Firstly, all `M` CIGAR operations are converted to `ID`, an insertion followed by a deletion. This change is shown in Figure 7, supplementing the original red alignment path with light gray cells. Next, computation proceeds one anti-diagonal row of width 2*b* + 1 at a time, centered on the alignment path. The computation of anti-diagonal rows shifts either right or downward at each step, governed by the previous CIGAR operation, `D` or `I`. These antidiagonal rows can be stored efficiently in matrix format, as demonstrated in Figure 7b. Transforming the banded |*r*|×|*r*| matrix *A* to a (2*b* +1) × 2 |*r*| matrix *B* saves significant space because nanopore sequencing read lengths |*r*| can be up to several million bases [8], while realignment works well with a band width of *b* = 30.

Offset arrays `INSs` and `DELs` can be precomputed using the CIGAR (Figure 7). Given a cell in matrix *B* with indices *i, j*, its position in matrix *A* can be computed using the following formula:

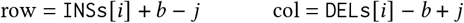

### 3.9 Time and Space Complexity

#### 3.9.1 Reference annotations

The worst-case time complexity for computing the reference annotations is 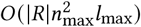, where |*R*| is the length of the reference *R*, and *n*_max_ is the maximum *n*-polymer considered, and *l*_max_ is the maximum *n*-polymer length. Since our *n*-polymer score matrix *N* is of size (6, 100, 100), *n*_max_ = 6, and *l*_max_ = 100. Thus, the time complexity is effectively *O* (|*R*|). Furthermore, these annotations must only be computed once, and cost can be amortized over all the reads that are aligned to the reference. We found the time required for reference annotations to be insignificant compared to alignment. These annotations require *O* (|*R* |*n*_max)_ space.

#### 3.9.2 Read alignment

Once the reference annotations and score matrices have been computed, the nPoRe algorithm requires *O(|R||r|)* time for each read *r*. The only additional overhead nPoRe incurs over Needleman-Wunsch with affine gaps is computing five Dynamic Programming (DP) matrices instead of three, as well as computing the new cell dependencies. All new penalties are conditional *O*(1) lookups. As discussed earlier in Section 3.8, follow-banding further reduces both the time and space complexity of alignment from *O* (|*R*||*r* |) to *O* (*b* |*r* |), where *b* is the band width. In total, the cost of aligning all reads is 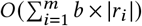, where *m* is the number of reads, or equivalently *O* (*db* |*R*|), where *d* is the average depth of coverage.

## 4 METHODOLOGY

All our code is open source and readily available at: https://github.com/TimD1/nPoRe.

### 4.1 Datasets

#### 4.1.2 Reference

We used the GrCh38 reference from the Genome-In-A-Bottle (GIAB) consortium [10].

#### 4.1.2 Reads

We obtained our FASTQ files from *ONT Open Datasets’* May 2021 re-basecalling of HG002 PromethION R9.4.1 data using Guppy 5.0.6. Specifically, we used flow cell `PAG07162`, prepared using the Short Read Eliminator (SRE) protocol [28]. Depth of coverage was approximately 60×. For training, we used `chr1-chr19`, and for testing we used `chr20-chr22`.

#### 4.1.3 Stratification Regions

Stratification BED regions were calculated for *n* = 1…*n*_max_ using the definition of *n*-polymers provided in Section 3.2. Regions were extended by a single base on each side (slop=1) to include variants occurring at the edges of *n*-polymer regions. These BEDS were then merged and complemented as necessary to create stratification BEDS for all repeat regions, tandem repeat regions, and non-repeat regions.

### 4.2 Pipeline

The full training and evaluation pipelines for all three Clair3 configurations tested are shown in Table 2. We used `minimap2` version 2.17-r954-dirty [12], clair3 version v0.1-r9 [31], whatshap version 1.0 [16], and `hap.py` version v0.3.14 [10]. All three variant callers were trained from scratch using our 60× HG002 dataset on MiniMap2-aligned reads for `chr1-chr19`, and tested on `chr20-chr22`. We first extended the retrained Clair3 baseline (`clair3`), phasing the input reads by haplotype and training a phased pileup candidate caller (`clair3-hap`). This was done because it significantly improves read concordancy, and leaving reads unphased when calling difficult single-haplotype variants might overshadow concordancy improvements gained by nPoRe’s alignment algorithm. This second baseline enables us the clearly delineate the gains from haplotype phasing and our nPoRe alignment algorithm. The final configuration was `clair3-npore-hap`, in which we performed ordinary variant calling with `clair3`, phased reads by haplotype, and then realigned them with nPoRe prior to variant calling.

**Table 2:** Clair3 training and evaluation pipelines.

| Step | clair3 | clair3-hap | clair3-npore-hap | Program |
| --- | --- | --- | --- | --- |
| align reads | ✓ | ✓ | ✓ | minimap2 |
| generate tensors | ✓ | ✓ | ✓ | clair3/CTP.py |
| train clair3 | ✓ | ✓ | ✓ | clair3/Train.py |
| call variants | ✓ | ✓ | ✓ | clair3/run_clair3.sh |
| phase variants |  | ✓ | ✓ | whatshap phase |
| phase reads |  | ✓ | ✓ | whatshap haplotag |
| phase truth VCF |  | ✓ | ✓ | whatshap phase |
| std truth VCF |  |  | ✓ | nPoRe/standardize_vcf.py |
| realign reads |  |  | ✓ | nPoRe/realign.py |
| gen hap tensors |  | ✓ | ✓ | clair3-hap/CTPHaps.py |
| train model |  | ✓ | ✓ | clair3-hap/Train.py |
| call variants |  | ✓ | ✓ | clair3-hap/run_clair3.sh |
| evaluate variants | ✓ | ✓ | ✓ | hap.py |

### 4.3 Haplotype Phasing

In order to add haplotype phasing information to Clair3, a single iteration of the ordinary pileup-based pipeline was first run. Proposed variants were then phased using `whatshap phase`, and reads were tagged by haplotype using `whatshap haplotag`. We then sorted reads by haplotype into three separate BAM files. When generating the input pileup tensor for training `clair3-hap` and `clair3-npore-hap`, for each position a pileup tensor was generated for both unphased reads, reads from the first haplotype, and reads from the second haplotype. These three pileup tensors were then concatenated to create a new input tensor for Clair3.

### 4.4 Truth VCF Standardization

As shown in Figure 1, our nPoRe realignment algorithm occasionally chooses a different variant representation than the original ground-truth VCF. nPoRe is comparatively more likely to call *n*-polymer INDELs than SNPs. We found that `clair3-npore-hap` variant calling performance suffers if we train Clair3 with the original ground-truth VCF, since the reads tend to report the same variant using a different representation. To mitigate this, we altered the ground-truth VCF so that it reports variants using the same representation our aligner tends towards.

To achieve this, we copied our reference FASTA to create two haplotype FASTAs, and applied the phased ground-truth variants to each haplotype FASTA, storing the new CIGAR. Using a mapping position of 0, the generated haplotype references, and associated CIGARs, we considered these ground-truth haplotype references to be reads and aligned them to the original reference using nPoRe. Any substitutions, insertions, or deletions in the resulting alignment were then parsed into a new standardized ground-truth VCF file. This process ensures that the new “standardized” truth VCF contains the same exact ground-truth sequence as the original VCF when applied to the reference FASTA, but reports variants in a manner consistent with nPoRe.

## 5 RESULTS

Figure 8 shows the calculated score matrices for 1- and 3-polymers, corresponding to the confusion matrices in Figure 3. In general, *n*-polymer INDELs are penalized less than the general-case affine gap INDEL penalty. Additionally, insertions are more common than deletions, and INDELs are more common in *n*-polymers of shorter repeat unit length (*n*).

**Figure 8:**
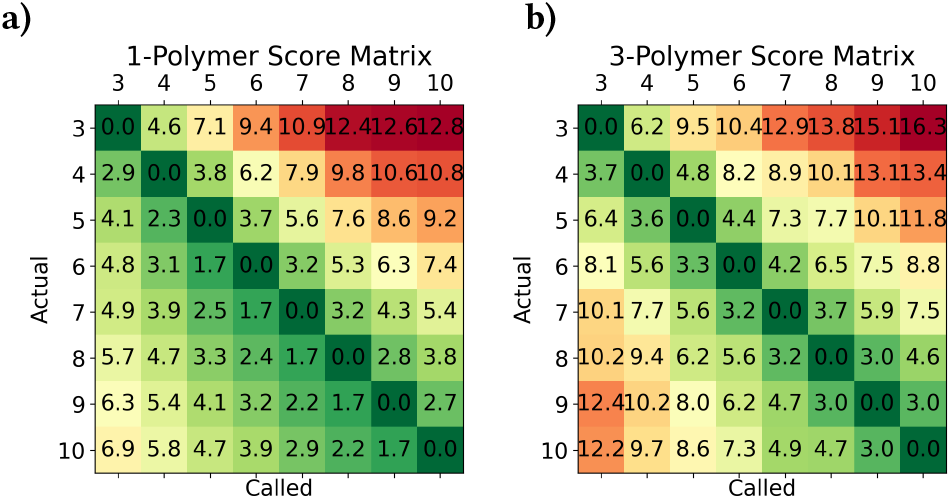
a) 1- and b) 3-polymer score matrices.

Read concordance, evaluated by Gini purity, is shown in Figure 9. Note that for the lower two plots, the y-scale is logarithmic. Gini pu-rity is defined as 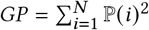, where *N* is the number of classes and ℙ(*i*) is the probability of class *i*. For the left two histograms in Figure 9, the classes are *A, C, G, T*, −, where −represents a deletion. For the right two histograms, the classes are all insertions between base *n* and *n* + 1, e.g. −, *A, AA, AAA, AT, ATT*. Since insertions do not directly align to a reference base, and the variable number of classes greatly affects the Gini purity score distribution, we chose to plot it separately. For insertions, there appears to be an approximately 10% increase for all non-perfect Gini purity scores. This is probably due to nPoRe’s increased likelihood of calling INDELs. For reference-aligned pileup bases, however, there is a marked ≈ 50% decrease in positions with Gini purity less than 0.5, demonstrating that nPoRe greatly improves read concordance in difficult regions.

**Figure 9:**
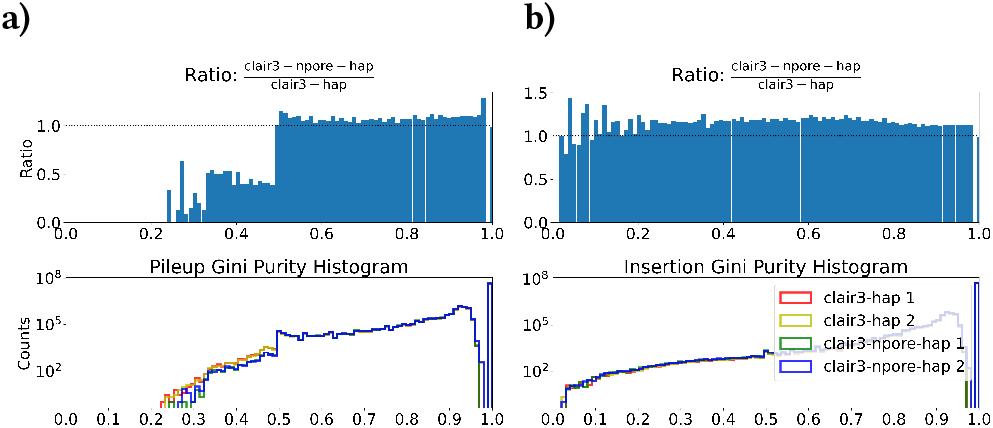
Read concordance: Gini purity histograms for a) pileup columns and b) insertions.

We performed our evaluations on a system with 2 × Intel Xeon E5 2697v3 2600MHz CPUs and 64GB total RAM. Although our nPoRe realigner accounted for 79.6% of total CPU time, it only accounted for 27.8% of the real runtime. Timing results are shown in Table 3. nPoRe’s CPU time was 51.6 × its real time on our system with 56 total cores, demonstrating that we took full advantage of the available parallelism.

**Table 3:** Timing Results.

| Step | Real Time | CPU Time |
| --- | --- | --- |
| align reads | 1,218:01 | 7,808:24 |
| generate tensors | 44:29 | 301:53 |
| train clair3 | 38:18 | 54:22 |
| call variants | 86:26 | 2,392:17 |
| phase variants | 347:58 | 344:05 |
| phase reads | 240:32 | 228:03 |
| index BAM | 1,138:09 | 1,750:13 |
| phase truth VCF | 364:23 | 360:03 |
| std truth VCF | 546:31 | 3,756:46 |
| realign reads | 2,168:15 | 111,792:55 |
| gen hap tensors | 1,363:23 | 3,162:48 |
| train model | 41:40 | 66:29 |
| call variants | 200:08 | 8,343:01 |

Figure 10 reports the performance of all three evaluated Clair3 pipelines, with precision and recall for SNPs and INDELs given separately for each sub-region. Results are reported for both the original (denoted +) and standardized (denoted •) ground-truth VCFs (see Section 4.4). The total fraction of SNPs/INDELs within each sub-region according to each truth VCF is included at the top of each plot.

**Figure 10:**
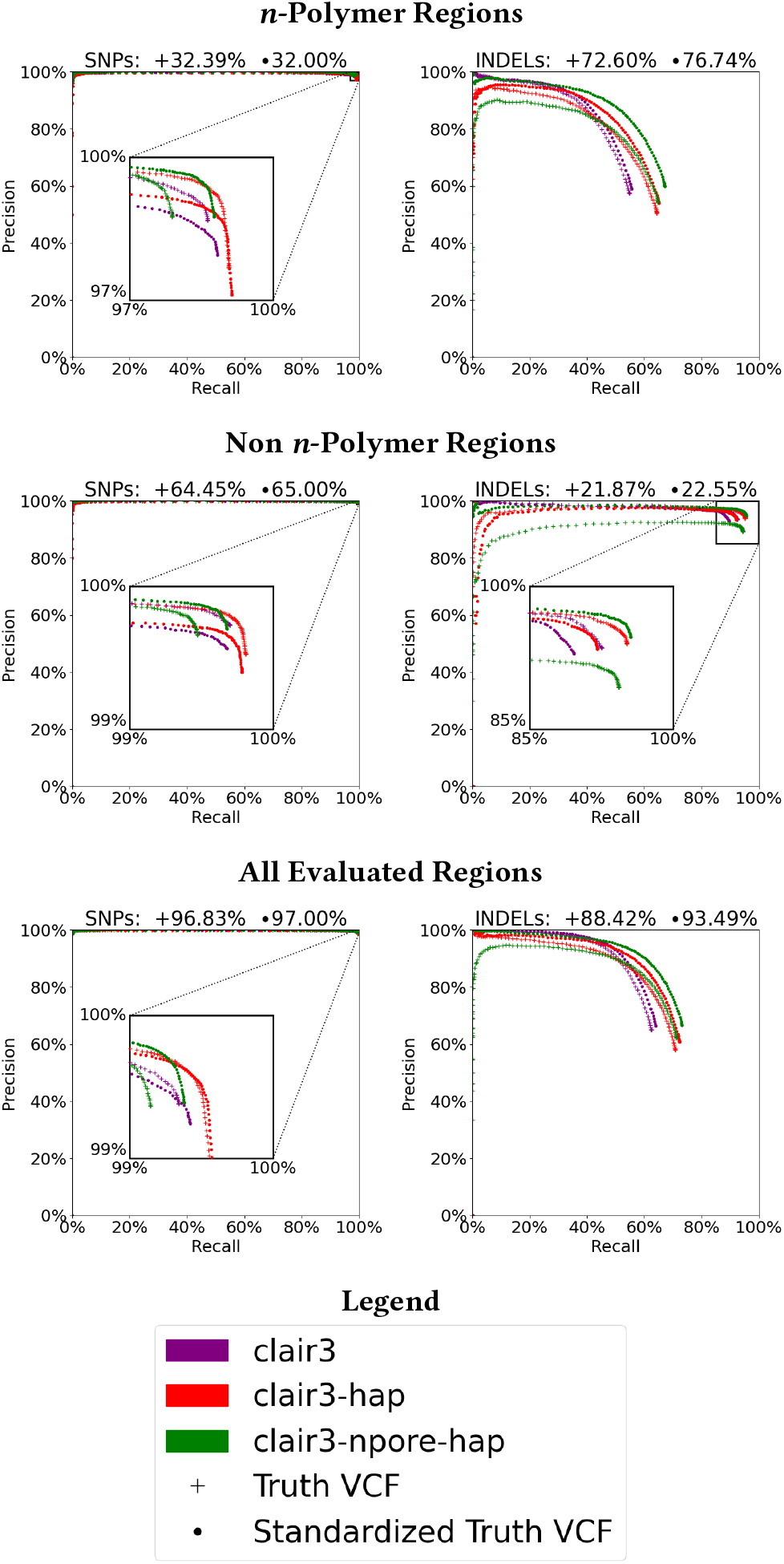
Accuracy results stratified by region.

We chose to perform evaluations using both VCFs because although they contain the exact same information, the “standardized” VCF was more likely to report several INDELs instead of several SNPs (due to the lower *n*-polymer shortening/lengthening penalty), and occasionally broke an INDEL up into several smaller INDELs. As a result, the standardized VCF had 18.05% more INDELs (31,104) and 1.45% fewer SNPs (155,163) than the original VCF (25,500 INDELs and 157,454 SNPs). `hap.py`’s `vcfeval` engine assigned partial credit for SNPs less frequently to nPoRe-aligned reads, and the standardized VCF resulted in apparent higher INDEL recall for all variant callers, due to the increase in total INDELs.

Figure 10 shows that *n*-polymer regions are responsible for the majority of INDEL errors, since with these regions excluded, INDEL precision and recall are both above 95%. Performance in tandem repeat regions alone is relatively good, and homopolymers account for the majority of remaining errors. For a fixed INDEL precision of 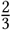, sorting reads by haplotype (`clair3→clair3-hap`) improves INDEL recall from 63.76% to 70.66%. Realigning reads with nPoRe (`clair3-hap→clair3-npore-hap`) further improves INDEL recall to 73.04%.

## 6 CONCLUSION

We identify the main source of nanopore germline small variant calling errors to be copy number variant false negatives in *n*-polymer regions, and show that context-agnostic affine gap penalties do not accurately reflect the likelihood of nanopore sequencing errors. To improve nanopore pileup-based variant calling accuracy, we explore correcting fine-grained read alignment.

This work extends Needleman-Wunsch affine gap alignment to include repeat-aware gap penalties for *n*-polymers. In doing so, we also develop “follow-banding” for efficient long read realignment and a method for standardizing ground-truth VCFs.

Our current nPoRe algorithm requires exact matches for consecutive repeat units within an *n*-polymer. A more lenient definition of *n*-polymers may result in broader application of this INDEL gap penalty and improve results further. nPoRe focuses on accuracy of alignment at the expense of efficiency and speed. As such, our Python implementation is relatively slow, despite our extensive usage of Cython and`multiprocessing`. Significant performance improvements can likely be gained by taking advantage of SIMD processing and replacing our large lookup tables (which have little spatial locality) with computation to improve cache efficiency. Nevertheless, we effectively demonstrate that read realignment improves read concordance and variant calling accuracy, and release nPoRe as an open source tool:

## ACKNOWLEDGMENTS

This project was supported by the National Science Foundation Graduate Research Fellowship under Grant 1841052. Any opinion, findings, and conclusions or recommendations expressed in this material are those of the authors and do not necessarily reflect the views of the National Science Foundation.

